# New adaptive peaks for crops – an example from improvement of drought-resilience of sorghum in Ethiopia

**DOI:** 10.1101/2021.10.18.464815

**Authors:** Techale Birhan, Hongxu Dong, Nezif Abajebel, Misganu Wakjira, Cornelia Lemke, Vincent Vadez, Andrew H. Paterson, Kassahun Bantte

**Affiliations:** Department of Horticulture and Plant Science, Jimma University, Ethiopia; Plant Genome Mapping Laboratory, University of Georgia, Athens, GA, USA; ICRISAT, Patancheru 502324, Andhra Pradesh, India; Department of Plant and Soil Sciences, Mississippi State University, Starkville, MS, USA

**Keywords:** adaptive traits, food security, genome-wide association studies, joint linkage analysis, sorghum, sub-Saharan

## Abstract

As the center of diversity for sorghum [*Sorghum bicolor* (L.) Moench], elite cultivars selected in Ethiopia are of central importance to sub-Saharan food security. Despite being presumably well adapted to their center of diversity, elite Ethiopian sorghums nonetheless experience constraints to productivity, for example associated with shifting rainfall patterns associated with climate change.
A sorghum backcross nested association mapping (BC-NAM) population developed by crossing thirteen diverse lines pre-identified to have various drought resilience mechanisms, with an Ethiopian elite cultivar, Teshale, was tested under three rain-fed environments in Ethiopia.
27, 15, and 15 QTLs with predominantly small additive effects were identified for days to flowering, days to maturity, and plant height, respectively. Many associations detected in this study corresponded closely to known or candidate genes or previously mapped QTLs, supporting their validity. Field tests show drought resilience to be improved by incorporation of adaptations from the diverse donor lines.
The expectation that genotypes such as Teshale from near the center of diversity tend to have a history of strong balancing selection, with novel variations more likely to persist in small marginal populations, was strongly supported in that for these three traits, nearly equal numbers of alleles from the donor lines conferred increases and decreases in phenotype relative to the Teshale allele. Such rich variation provides a foundation for selection to traverse a ‘valley’ of reduced yield and arrive at a new ‘adaptive peak’, exemplifying the nature of efforts that may be necessary to adapt many crops to new climate extremes.

**Societal Impact Statement:** In Ethiopia, agriculture is the largest economic sector and contributes 48% of the nation’s GDP, and sorghum provides more than one third of the cereal diet and is widely grown for food, feed, brewing, and construction purposes. With a worldwide water crisis looming, developing drought tolerant sorghum is vital in rain-fed environments, particularly in sub-Saharan Africa. Addressing such issues often requires a far-reaching approach to identify and incorporate new traits into a gene pool, followed by a period of selection to re-establish an overall adaptive phenotype. This study revealed that with the enormous altitudinal variation of a country such as Ethiopia, somewhat different lines may be needed for different locales.

## Introduction

Sorghum [*Sorghum bicolor* (L.) Moench], a short day C_4_ tropical grass native to Africa, is exceptional in its wide range of adaptation. Since its primary domestication near present-day Sudan approximately 5,000 years ago (Winchell *et al*. 2017), sorghum has been introduced to diverse climates across Africa, Asia, and the Americas (De Wet and Harlan 1971). Flowering time plays a central role in plant adaptation to local environmental conditions, with local ideotypes ranging from short-day forms for the tropics to day-neutral rapidly flowering forms for high temperate latitudes with short growing seasons. Genetic improvement of sorghum and other cereal crops has also adjusted plant stature to meet needs ranging from provision of extensive biomass forage and thatch for building (Blümmel & Rao 2006; Mathur *et al*. 2017; Murray *et al*. 2008; Tesso *et al*. 2008), to dwarf forms ideal for mechanical harvesting and to avoid lodging and other natural hazards. Indeed, the latter is exemplified by the success of the Green Revolution, in which grain yield increased substantially in rice (*Oryza sativa* L.) and wheat (*Triticum aestivum* L.) by the introduction of semi-dwarf traits (Hedden 2003; Peng *et al*. 1999). Similarly, a Sorghum Conversion Program backcrossed genomic regions conferring early flowering and dwarfing from an elite donor into approximately 800 exotic sorghum accessions, advancing adaptation to grain and biomass production in temperate regions (Stephens *et al*. 1967).

Flowering time and plant height are quantitative in nature. Cultivated grain sorghum varieties typically flower between 45 and 120 days after planting under various day lengths, and could range from 2 to 18 feet in height depending on the dwarfing genes they contain. Classical studies suggested that sorghum flowering and plant height are each controlled by at least four major loci (*Ma1-4* and *Dw1-4*, respectively) (Quinby 1974). Additional maturity loci (*Ma5*-*6*) with large effects were reported by Rooney and Aydin (1999). Among the six major maturity genes, Cuevas et al. (2016) revealed a phosphatidylethanolamine-binding (PEBP) protein, Sb06g012260, to be the *Ma1* gene, as supported by independent lines of evidence including fine mapping, association genetics, mutant complementation, and evolutionary analysis. Others have suggested positional candidates (not confirmed by transformation) for sorghum flowering genes including a pseudo-response regulator protein (PRR37) for *Ma1* (Murphy *et al*. 2011); a SET and MYND (SYMD) domain lysine methyltransferase for *Ma2* (Casto *et al*. 2019); phytochrome B and phytochrome C for *Ma3* and *Ma5*, respectively (Yang *et al*. 2014); and *SbGHD7*, a repressor of flowering in long days, for *Ma6* (Murphy *et al*. 2014). Among the four ‘major’ sorghum dwarfing genes, *Dw3* encodes an auxin efflux carrier, PGP19 (Multani *et al*. 2003), while candidate genes unconfirmed by complementation include a putative membrane protein that possibly involve in brassinosteroid signaling for *Dw1* (Hirano *et al*. 2017; Yamaguchi *et al*. 2016) and a protein kinase for *Dw2* (Hilley *et al*. 2017).

The control of flowering and height are much more complex than suggested by classical genetics, as genetic linkage analyses have revealed numerous additional loci (Zhang *et al*. 2015), among which some show major effects under various genetic backgrounds. A new recessive dwarf mutation, *dw5*, was recently isolated from a mutagenized BTx623 mutant library, but its molecular function is yet to be studied (Chen *et al*. 2019).

In natural populations, genotypes at the center of diversity tend to be under strong balancing selection, with novel variations more likely to persist in small marginal populations conducive to intense selection and/or fixation by drift. Here, a genotype selected in and presumably well adapted to its center of diversity is crossed to each of thirteen diverse lines from across the natural and introduced range of sorghum, chosen for their increased capacity to extract water from the soil or for their exceptional transpiration efficiency (Vadez *et al*. 2011), but also with diverse morphology and growth habit. A backcross nested association mapping (BC-NAM) population such as was produced here, consisting of multiple families crossed to a common tester, allows one to catalogue allelic variants at numerous QTLs and determine their contribution to phenotype and distribution across diverse germplasm (Yu *et al*. 2008). We evaluated this BC-NAM population under three natural environments that are prone to drought in Ethiopia, which afforded the opportunity to study adaptive traits under rain-fed environments. Joint-linkage and GWAS approaches were applied to map the genetic basis of flowering time, maturity, and plant height.

## Materials and Methods

### Plant materials and population development

The 13 founder lines used for the population development were obtained from ICRISAT (Table 1). These 13 diverse founder lines were selected based on their diverse spectrum of drought responsiveness traits, especially in their water extraction ability and transpiration efficiency (Vadez *et al*. 2011). IS2205, IS14446, and IS23988 were characterized by excellent water extraction ability; IS3583, IS14556, IS16044, IS16173, IS22325, IS10876, and IS15428 showed good transpiration efficiency; IS9911, IS14298, and IS32234 showed good harvest index (Vadez *et al*. 2011). The recurrent common parent, Teshale, is an Ethiopian variety of caudatum origin which is highly preferred by the Ethiopian farmer for its grain quality and high yield. Seeds of Teshale were sourced from Melkassa Agricultural Research Center in Ethiopia. The sorghum BC-NAM population was developed using a nested design, crossing the common parent, Teshale, with the selected founder line and backcrossing the resulting F_1_ to Teshale to produce BC_1_F_1_ families. Crossing was done by hand emasculation of normal bisexual florets (approximately 50 per panicle) of Teshale, transferring pollen from the founder lines to the stigma of the emasculated florets 3-5 days later. Following the BC_1_F_1_ generation, genotypes were continuously selfed to BC_1_F_4_ via single seed descent. Finally, the BC_1_F_4_ generation was used for genotyping and phenotyping. The populations were developed between 2013/14 to 2016/17. Below, when referring to an individual BC_1_F_4_ population, we will use the name of the alternate parent (*e.g*., the IS9911 population; Table 1).

**Table 1.** Description of the sorghum BC-NAM population

| Donor<br>Name□ | Country of<br>origin§ | Genetic | No. of SNPs | Average | Pop. size<br>in Kobo | Pop. | Pop. size |
| --- | --- | --- | --- | --- | --- | --- | --- |
|  |  | similarity<br>with<br>Teshale |  | percentage<br>of Teshale<br>genome |  | size in<br>Meiso | in<br>Sheraro□ |
| IS10876 | Nigeria | 0.610 | 2592 | 75.0% | 135 | 151 | 153 |
| IS14298 | South Africa | 0.752 | 2809 | 70.7% | 121 | 132 | 131 |
| IS14446 | Sudan | 0.776 | 2425 | 72.5% | 134 | 145 | 149 |
| IS14556 | Ethiopia | 0.896 | 1888 | 88.5% | 32 | 34 |  |
| IS15428 | Cameroon | 0.811 | 2963 | 79.9% | 124 | 142 | 144 |
| IS16044 | Cameroon | 0.723 | 1597 | 78.5% | 36 | 37 |  |
| IS16173 | Cameroon | 0.706 | 1541 | 76.5% | 98 | 111 | 112 |
| IS2205 | India | 0.884 | 2283 | 75.5% | 36 | 40 |  |
| IS22325 | Botswana | 0.753 | 2494 | 68.9% | 101 | 120 | 120 |
| IS23988 | Yemen | 0.770 | 2019 | 83.6% | 34 | 42 | 36 |
| IS32234 | Yemen | 0.854 |  |  |  |  |  |
| IS3583 | Sudan | 0.796 | 2285 | 83.8% | 104 | 119 | 119 |
| IS9911 | Sudan | 0.645 | 1932 | 78.3% | 76 | 82 | 82 |

□ Founder lines used in the BC-NAM population. Note that population IS32234 was discarded due to severe contamination.
§ Source information is obtained from http://genebank.icrisat.org/IND/Passport?Crop=Sorghum.
□ Populations IS2205, IS14556, IS16044, IS32234 were not included in Sheraro due to space constraint.

### Experimental design and trial management

Parental lines and BC_1_F_4_ lines were initially evaluated in 2017 at two environments in Ethiopia: Kobo (12°09’N, 39°38’E) and Meiso (09°14’N, 40°45’E). Due to strong moisture stress during the sowing season (July 2017), large numbers of seeds failed to germinate at Kobo and Meiso fields, resulting in uneven planting density at these two locations. Therefore, a third field trial was arranged at Sheraro (14°23’N, 37°46’E) in 2018. These three locations represent major sorghum production regions in Ethiopia and were selected for their natural drought conditions (Figure 1). Irrigation was not available at these three locations and thus this BC-NAM population was challenged with rain-fed condition. An alpha lattice design with two replications was used at each location. Seeds of each line were sown into one-row plots, with 75 cm between rows for a net plot size of 0.75 m × 4 m. Inorganic fertilizers DAP and Urea were added at the rates of 100 and 50 kg ha^−1^ as side dressing during sowing and three weeks after sowing, respectively. Thinning of seedlings was done three weeks after sowing, to 20 cm spacing between individual plants. Therefore, individual plots without plant loss would consist of 20 plants. Weeding and other cultural practices were carried out as needed.

**Figure 1.**
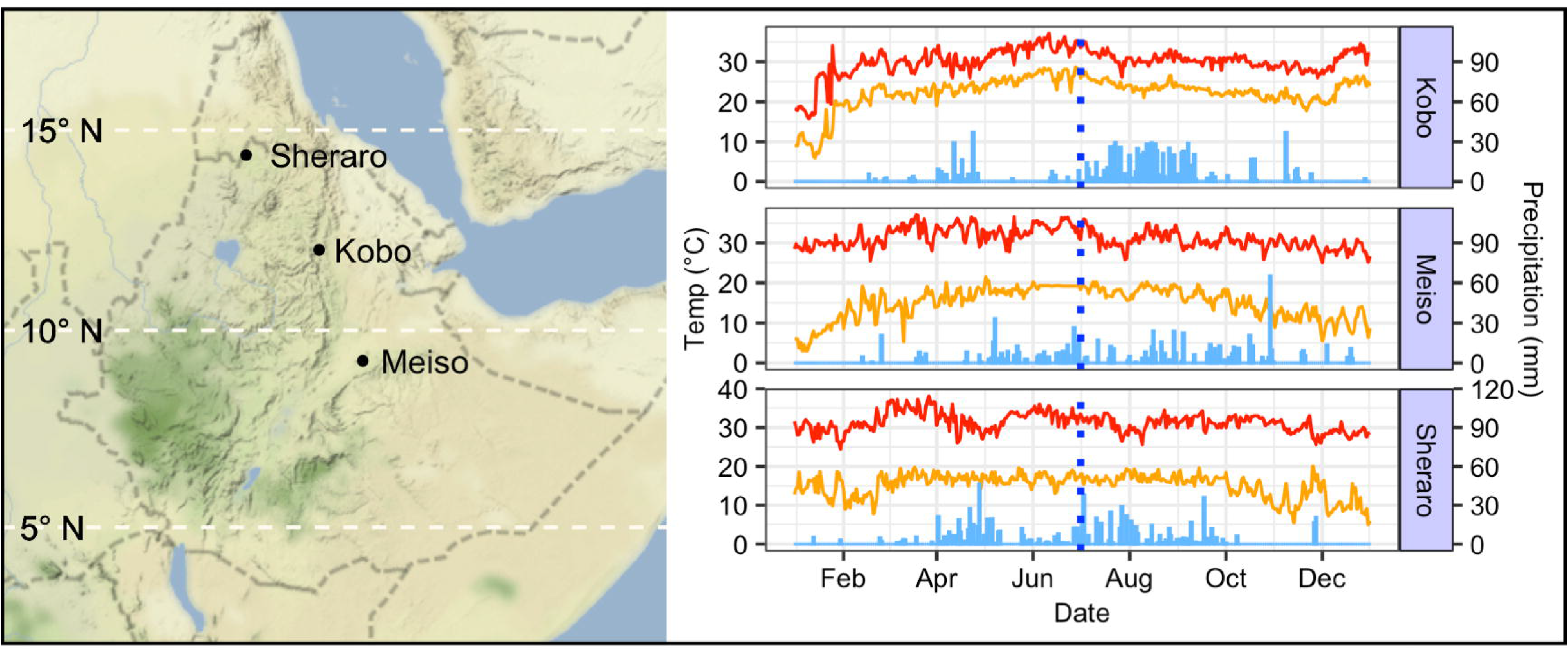
Geographic location and environmental condition of the three field trials in Ethiopia. Red lines and orange lines represent daily maximum and minimum temperatures, respectively. Vertical solid blue bars represent daily precipitation, and vertical dashed lines represent the sowing time in July.

Flowering and plant height traits were evaluated in this BC-NAM population. Days to flowering (DF) was defined as the number of days until 50% of plants per plot were in anthesis. Days to maturity (DM) was the number of days until 50% of plants per plot reached physiological maturity. Plant height (PH) was the mean of five representative plants per row, measured from the base to the tip of panicle after physiological maturity in centimeters.

### Phenotypic data analysis

Analysis of variance (ANOVA) was first conducted across all three environments to test significance of environment, family, genotype nested within family, family by environment interaction, and genotype nested within family by environment interaction. We compiled weather data including daily precipitation, minimum and maximum temperature from nearby weather stations during the calendar years of field trials (Figure 1). The cumulative precipitation before sowing (Jan-June) were 194.4 mm and 313.6 mm at Kobo and Meiso, respectively, whereas it was 401.2 mm at Sheraro. The lower soil moisture at Kobo and Meiso likely explained why many seeds failed to germinate compared to Sheraro. Given the distinct conditions across these three environments (Figure 1), trait best linear unbiased predictions (BLUPs) were estimated for each line within each environment using a mixed linear model implemented in the lme4 package (Bates *et al*. 2015). All model terms were treated as random effects except for grand mean in the following model:

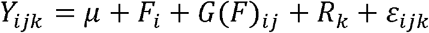

where *Y* represents raw phenotypic data, *μ* is grand mean, *F* is the individual BC-NAM family, *G*(*F*) is genotype nested within family, *R* is replication, and *ε* is random error. Pearson correlation coefficients between traits were calculated using line BLUPs. Broad-sense heritability was determined as the proportion of total phenotypic variance explained by the combined family and line terms using the equation:

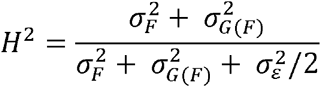

where 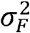 is the variance explained by family term, 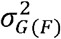 is the variance explained by individual lines, and 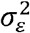 is the random error variance.

### Marker development and genomic analyses

Genomic DNA was extracted from freeze-dried leaves using a CTAB (cetyltrimethylammonium bromide) protocol (Doyle & Doyle 1987). Twelve randomly selected DNA samples from each population were checked for genomic integrity on 2% agarose gels before library construction. DNA concentration of each sample was quantified using a Qubit Fluorometer dsDNA system (Invitrogen, Carlsbad, CA) and diluted to 20 ng/μl. Libraries were constructed using a *Pst*I-*Msp*I enzyme system (Poland *et al*. 2012) with modifications based on Clark *et al*. (2014). DNA samples were digested with the rare-cutting enzyme *Pst*I-HF (High-Fidelity; New England Biolabs Inc., Ipswich, MA, USA) and the common-cutting enzyme *Msp*I (New England Biolabs Inc., Ipswich, MA, USA), then ligated to a unique barcode adapter and a common adapter. A total of 192 samples (*i.e*. corresponding to 192 unique barcodes) were pooled in one library, and 200-500 bp fragments were extracted from a 2% agarose gel after electrophoresis and purified using a Qiagen Gel Extraction Kit (Qiagen, Hilden, Germany). The purified DNA was PCR amplified using 2× GoTaq Colorless Master Mix (Promega, Madison, WI, USA), and PCR product was extracted as above to eliminate primer-dimers. All GBS libraries were sequenced on a NextSeq500 (Illumina, San Diego, CA, USA) with 150 bp single-end reads at the University of Georgia Genomics and Bioinformatics Core. The 14 parents were replicated at least twice in order to improve coverage and to correctly call SNPs in progeny. SNP calling was performed with GBS v2 pipeline in TASSEL (Bradbury *et al*. 2007) using version 2.1 of the *Sorghum bicolor* genome (Paterson *et al*. 2009). To remove low-quality genotypic data, raw genotypes were filtered for tag coverage (tag found in >5% of taxa), minor allele frequency (MAF>0.03), and single marker missing data (<0.8). Trimming SNPs with 5% missing data and trimming nearby (<100 bp) SNPs with identical genotypes yielded 4395 SNPs for further analysis.

### Genomic analyses

Principal component analysis (PCA) was first performed within each of these 13 BC-NAM populations to identify putative contaminants. One to seven individuals within each population exhibited high levels of errant genotypes and could not cluster with their respective population; IS32234 was of very small size (N = 25) and grouped into two clusters (Figure S1), probably because of contamination, mistaken parental identity, or incorrect pollination. Therefore, 54 individuals (25 individuals of IS32234 and 29 putative contaminants from the other 12 populations) were excluded and 1171 BC_1_F_4_ lines were retained for all analyses. A composite PCA of the retained 1171 BC_1_F_4_ lines was then conducted. Recurrent parent allele frequencies, genome-wide heterozygosity, and SNP monomorphism rates were calculated in R with customized scripts. Linkage disequilibrium (LD) was calculated as squared allele frequency correlations (*r^2^*) in VCFtools (Danecek *et al*. 2011). Decay of LD with distance in base pairs between sites was modeled using the nonlinear regression model of Hill and Weir (1988). Polymorphic markers within each population were used to estimate the percentage of common parent genome present in each of the derived BC_1_F_4_ lines. Whole genome mean, maximum, and minimum percentages of the common parent genotype were estimated for each population.

### Marker-trait association

To map QTL in the BC-NAM population, we used a joint-linkage (JL) model (Buckler *et al*. 2009) and GWAS approach. In JL analysis, we only included eight large populations (N > 80, Table 1), removing IS14556, IS16044, IS2205, and IS23988 due to their small size. This decision was supported by the consideration that small population size would lead to reduced power in QTL detection, underestimation of QTL number, and overestimation of QTL effect (Vales *et al*. 2005; Yu *et al*. 2008). For JL mapping across the eight populations, a new SNP dataset was created to track parent-of-origin within each population. The common parent Teshale genotypes were set to 0, alternative parent genotypes were set to 1, and heterozygous genotypes were set to 0.5. Monomorphic SNPs within each family were set to missing, and missing data were imputed as the mean of the nearest flanking markers weighted by physical distance (Tian *et al*. 2011). Therefore, the result can be interpreted as the probability that a SNP comes from the donor parent, and adjacent SNPs are always in high linkage disequilibrium with each other in this dataset because it reflects only the meiosis that occurred during the creation of each BC_1_F_4_ population. Joint-linkage analyses were performed using the Stepwise Plugin of TASSEL 5 (Bradbury *et al*. 2007). SNP effects were nested within populations to reflect the potential for unique QTL allele effects within each population. Although it is unlikely that there is a unique allele for each population at every QTL, this nested model provides a statistical framework for modeling multiple alleles at any given QTL. Therefore, based on this model, multiple allelic effects, as opposed to only two, are reported for each QTL. Multi-parental mapping using the GWAS approach used the unmodified genotypic dataset of all 12 populations. In each approach, the population term was included as a fixed effect to account for the inherent structure in the BC-NAM lines. The critical difference between joint-linkage mapping and GWAS is that joint-linkage mapping relies on parent-of-origin information while GWAS only uses allele state information of markers.

All JL QTLs identified in this study were compared to the Sorghum QTL Atlas database described in Mace *et al*. (2019), which collated the projected locations of *c*. 6000 QTL or GWAS loci from 150 publications in sorghum from 1995 to present. QTL comparison was conducted based on their projected physical locations on version 2.1 of the *S. bicolor* genome. QTLs for the same trait were declared as common QTLs if they showed overlapping confidence intervals. Some QTLs from maps of low resolution occasionally span whole chromosomes and were not considered for comparison. In addition, sorghum genes containing or directly adjacent to SNP associations were searched using BEDOPS (Neph *et al*. 2012) and standard UNIX scripts.

### Data availability

Sequencing data are available in the NCBI Sequence Read Archive under BioProject ID PRJNA687679. Data analysis scripts have been deposited to GitHub (https://github.com/hxdong-genetics/Ethiopian-Sorghum-BC-NAM). Please contact the corresponding author for other data.

## Results

### Genetic diversity and structure of the BC-NAM population

To evaluate the genetic diversity and structure of the BC-NAM population, we characterized the 1171 BC_1_F_4_ lines at 4395 high-quality GBS SNPs, which corresponds to an average density of one SNP per 1.5 Mb. Among these 4395 SNPs, 3029 (68.9%) were located within genic regions (Dataset S2). Principal component analysis showed individuals of each population to be clearly clustered (Figure 2A). The first three principal components explained 7.7%, 5.3%, and 3.9% of the variance, respectively. The IS22325-derived population exhibited the most genetic difference with the other populations based on PC2, followed by IS14298 based on PC3 (Figure 2A). Genetic similarity between the common parent and each alternate parent was lowest with IS10876 (0.610) and highest with IS14556 (0.896), which led to variation in monomorphism across the genome within each population (Table 1).

**Figure 2.**
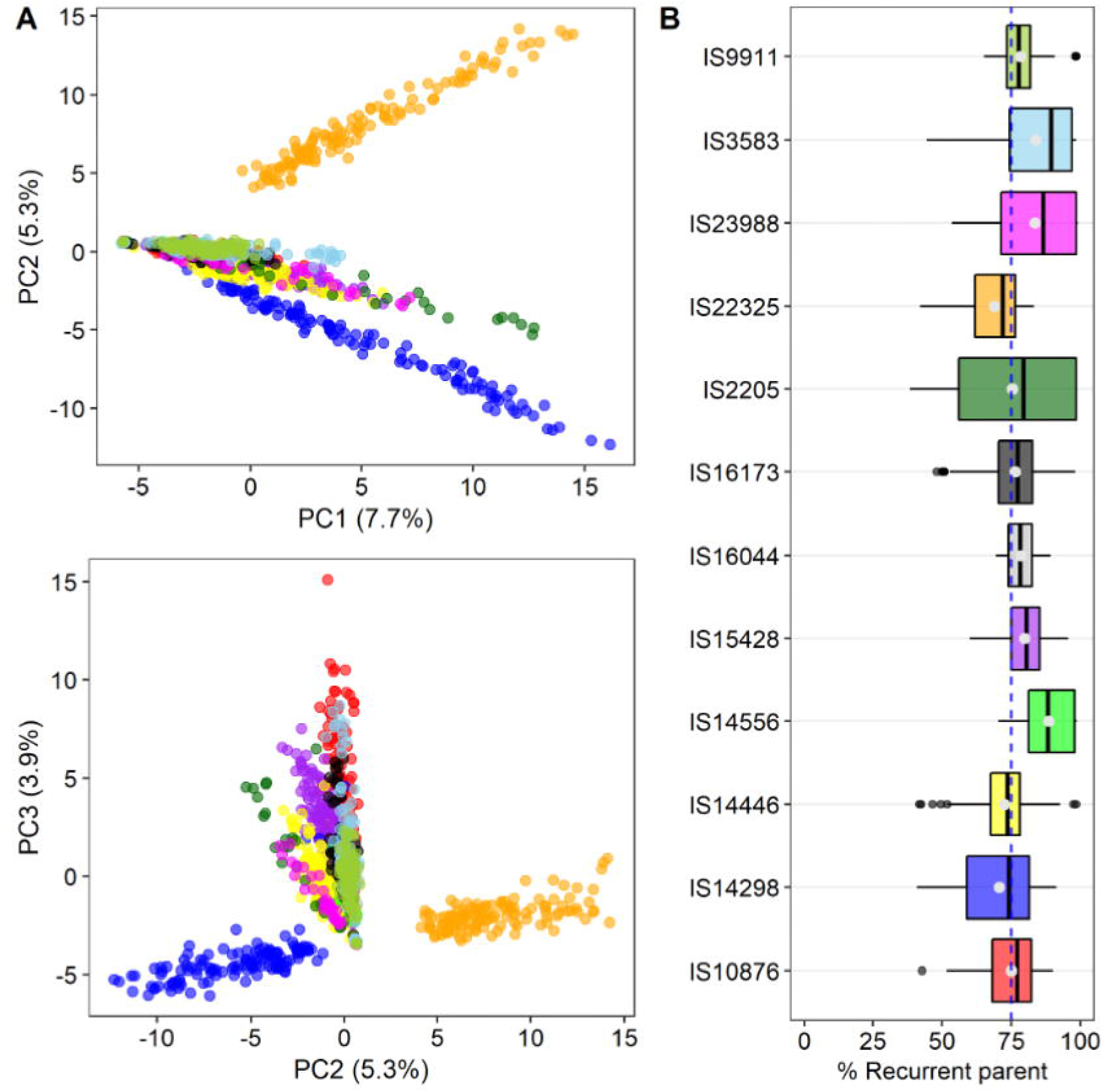
Genomic diversity of the BC-NAM population. (A) Principal component analysis across 1171 BC_1_F_4_ lines at 4395 SNPs. Variance explained by each PC was shown in parenthesis; (B) Boxplot distribution of the percentage of recurrent parent genome present in each population. The theoretical 75% value was indicated with a vertical dashed line. Dot inside each boxplot is mean, and the vertical line is median.

The overall mean percentage of recurrent parent genome (PRPG) was about 76.3% for all the populations but varied considerably between populations, from 68.9% in IS22325 to 88.5% in IS14556 (Figure 2B, Table 1). Three populations including IS22325 (68.9%), IS14298 (70.7%), and IS14446 (72.5%) showed lower mean PRPG than the theoretical 75% (Figure 2B, Table 1). The highest PRPG in IS14556 echoes its highest genetic similarity with the common parent (0.896, Table 1). Although we did not impose artificial selections during population development, the natural drought conditions in Ethiopia could have selected plants with local adaptation because some seedlings failed to germinate (K. Bantte, personal communications), and thereby explained higher than expected PRPG in the other ten populations (Figure 2B). Indeed, a common set of 58 BC-NAM lines lost one of two replicates at both Kobo and Sheraro (Table S1; i.e., presumably due to poor germination associated with moisture stress). The average PRPG in these 58 lines was 74%, compared to 76% in the remaining lines without severe plant loss (Dataset S3). This indicated that BC-NAM lines with poor germination enriched for exotic parent alleles. The minimum percentage of recurrent parent genome also varied between the populations from as low as 38.44% for one line from the IS22325 population to 70.33% for a line from the IS14556 population (Dataset S3). The theoretical range of PRPG after one generation of backcross without selection is 50-100%. Few individuals with PRPG lower than 50% were likely derived from F_1_ seeds rather than BC_1_s. These few individuals were expected to have minimal impact on association analyses given their overall consistent clustering within respective population (Figure 2A). The maximum percentages of recurrent parent genome varied from a high of 98.71% for a line from the IS22325 population to as low as 83.21% for a line from the IS22325 population (Figure 2B, Dataset S3).

Linkage disequilibrium (LD) decayed to 0.2 at *c*. 260 kb in this BC-NAM population (Figure 3A), larger than the 100-150 kb in sorghum diversity panels (Hamblin *et al*. 2005; Morris *et al*. 2013) due to the backcross breeding scheme. One generation of backcross recovers 75% of the recurrent parent genome, resulting in long haplotypes of the recurrent parent being maintained across the genome in these BC_1_F_4_ lines. LD is of great importance for the design of association studies to identify the genetic basis of complex traits. Given that the genome length of sorghum is *c*. 730 Mb (Paterson *et al*. 2009), a minimum of *c*. 2800 markers (730/0.26) would provide an average of one marker per LD block in the present study. Therefore, the 4395 SNP markers here are expected to sample most genetic variation in this BC-NAM population, with high power to detect marker-trait associations.

**Figure 3.**
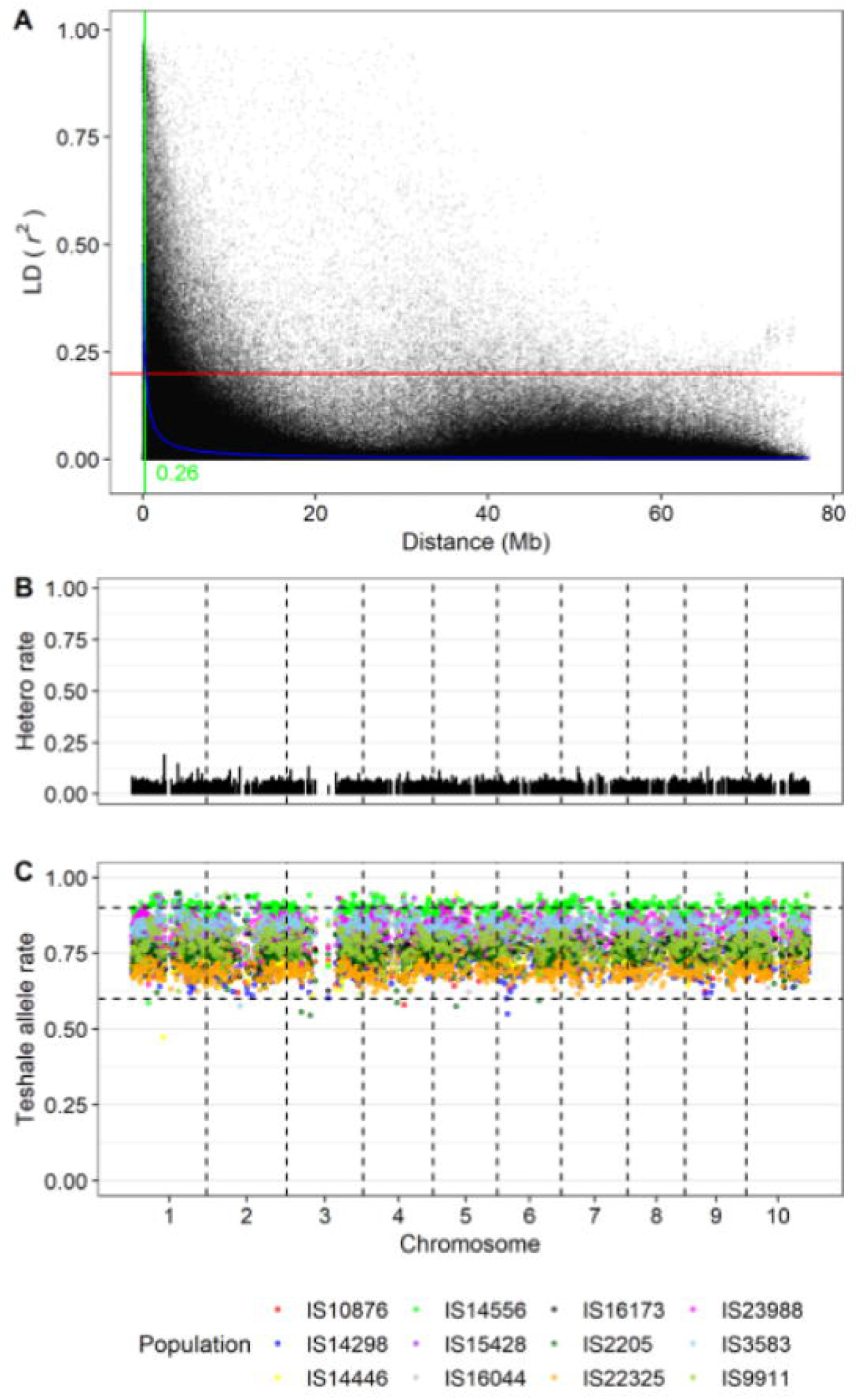
Genomic properties of the BC-NAM population. (A) Linkage disequilibrium (LD) decay plot, using non-linear model described in Hill and Weir (1988). Each dot represents pairwise *r*^2^ between SNPs within chromosome. (B) The percentages of BC-NAM lines with heterozygous genotypes (“Hetero rate”) across the 4395 SNPs. (C) Segregation of recurrent parent alleles. Each dot represents the recurrent parent allele frequency within a certain population, colored as shown in the legend. Horizontal dashed lines stand for 0.60 and 0.90 thresholds for significant segregation distortion.

The genetic structure and diversity of the BC-NAM population might have been affected by natural selection during population development, which can lead to decreased residual heterozygosity and segregation distortion. Heterozygosity rate in the BC-NAM population was 0.0606 (Figure 3B), slightly lower than the expected value for the BC_1_F_4_ generation (0.0625). The decreased heterozygosity also echoes the slightly higher percentage of recurrent parent genome (76% vs. 75%, Figure 2B). Across the genome 85.67% of markers exhibited heterozygosity <= 0.1 (Dataset S2), suggesting that natural selection had little effect overall. The frequency of alleles from the common parent, Teshale, was close to the neutral expectation (75%: Figure 3C), suggesting no overall selection for or against common parent alleles. Still, a small proportion of markers showed skewed segregation, for either the common parent (e.g. IS14556), or alternate parent (IS22325) allele (Figure 3C), suggesting selection at some loci. No clear difference was observed among families in terms of proportion of distorted markers, and skewed chromosome regions were generally specific to one or a few families.

### Phenotypic variation within and between families

We evaluated the BC-NAM population over three environments (Kobo, Meiso, and Sheraro) representing major sorghum cultivation zones in Ethiopia for days to flowering, days to maturity, and plant height (Figure 4). As shown in Figure 1, these three locations had similar temperature profiles, but different daily precipitation distributions. The cumulative precipitation before sowing (Jan-June) were 194.4 mm and 313.6 mm at Kobo and Meiso, respectively, whereas it was 401.2 mm at Sheraro. In particular, from May to June, Kobo only received 32.6 mm precipitation, while Meiso and Sheraro received 250.2 mm and 167.2 mm, respectively (Figure 1). As a result, severe plant losses occurred at Kobo and Meiso, with 362 and 195 individuals lost in one of the two replicates, respectively (Table S1). In Sheraro, virtually all plants survived -- only two individuals lost one replicate (Table S1). Given the precipitation data, plant losses at Kobo and Meiso were presumably caused by poor germination due to moisture stress (K. Bantte, personal communications). Multi-environment ANOVA confirmed strong environmental effect and G×E interactions (*P* < 2.2E-16; Table S2). Between G and G×E, mean squares of G were generally twice the magnitude of G×E for the three traits (Table S2). Thus, trait BLUPs were estimated within each environment and trait-marker association analyses were performed separately for each environment.

**Figure 4.**
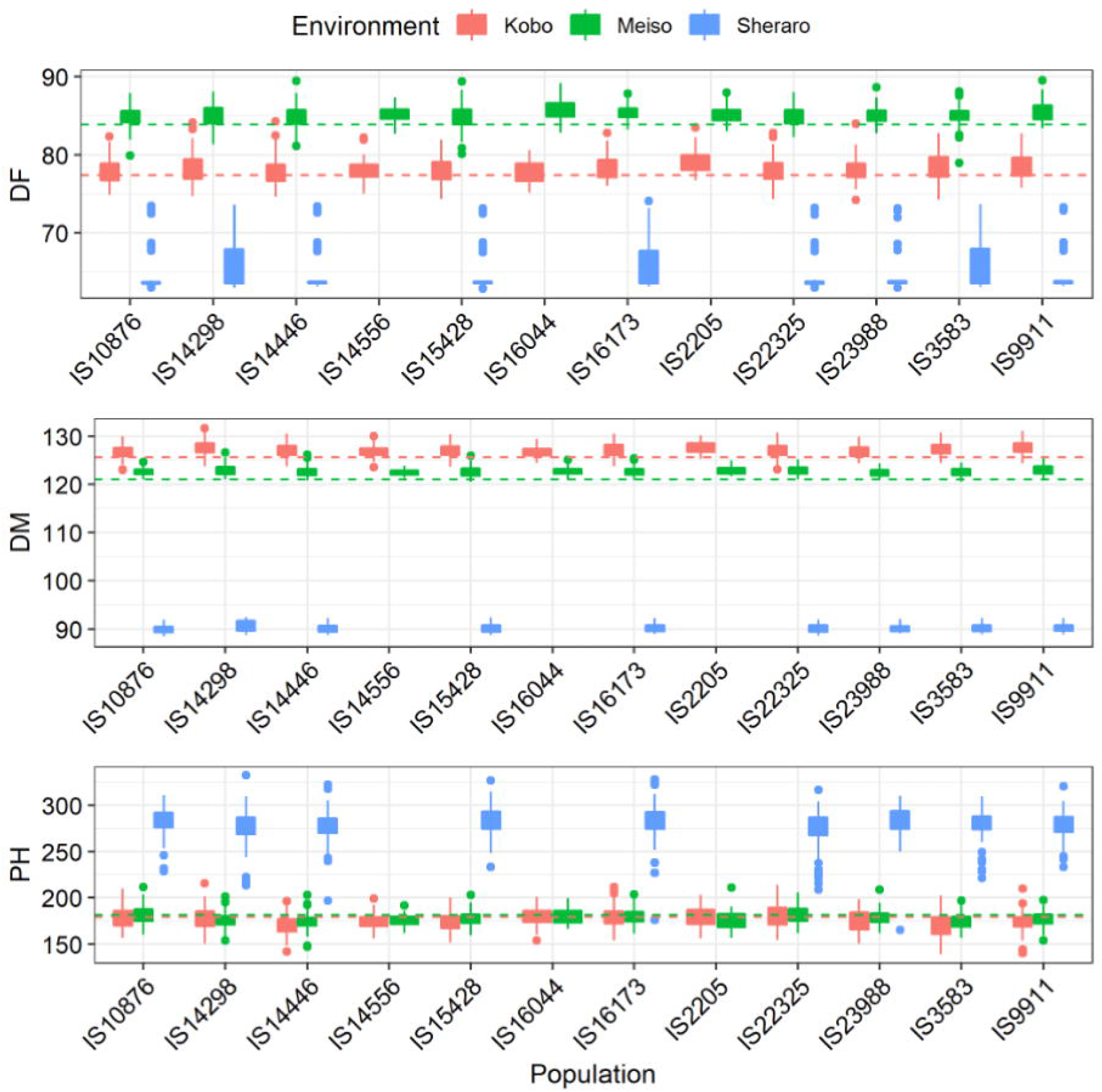
Phenotypic variation within each individual population at three environments (Kobo, Meiso, Sheraro). Phenotypes were shown for DF: days to 50% flowering, DM: days to maturity, PH: plant height. Inside each plot, the horizontal dashed lines represent trait mean of the common parent Teshale.

Plants in the least drought-stressed location, Sheraro, flowered and matured earliest while also being nearly twice the height of those at the other locations. Average flowering of the BC-NAM population occurred in Sheraro at 65 d after sowing, followed by Kobo at 78 d and Meiso at 85 d (Figure 4, Table S3), reaching maturity in Sheraro at 90 d, followed by Meiso at 123 d, and Kobo at 127 d. The BC-NAM populations were of similar average plant height at Kobo and Meiso, at 175.7 cm and 177.7 cm, respectively; but significantly taller at Sheraro with 279.6 cm. Trait distributions within each of the 12 populations at the three environments were similar, in terms of population mean and standard deviation (Figure 4, Dataset S1). Among the 12 donor parents and the recurrent parent Teshale, days to flowering ranged from 75.9 to 80.1 and 83.5 to 87.1 in Kobo and Meiso, respectively; and days to maturity ranged from 125.3 to 129.7 and 121.0 to 124.2 in Kobo and Meiso, respectively (Dataset S1). In contrast, plant height was more variable, parental lines ranging from 149.3 cm to 201.8 cm in Kobo and 156.5 cm to 202.6 cm in Meiso (Dataset S1). Due to the backcross breeding scheme, individual population means were generally close to those of the recurrent parent Teshale (Figure 4).

Broad-sense heritabilities of these three adaptive traits were generally higher in the least drought-stressed location, Sheraro (Table 2). Days to flowering heritability was high in Sheraro (0.71) but low in Kobo (0.33) and Meiso (0.30). Days to maturity heritabilities were consistently low across all three environments (0.25-0.34), which was probably due to the very limited phenotypic variance (Figure 4). Heritability of plant height followed a similar pattern as days to flowering, with the highest value observed in Sheraro (0.75), medium in Kobo (0.55), and relatively low in Meiso (0.39). Stress responses may have masked genetic potential, and/or invoked different and more complex genetic controls than in favorable environments (Paterson *et al*. 2003). Moreover, medium to high positive correlations were observed between days to flowering and days to maturity (Kobo: 0.69; Meiso: 0.59; Sheraro: 0.33), whereas negligible or negative correlations were found between plant height and days to flowering (Kobo: 0.12; Meiso: −0.28; Sheraro: −0.03) and between plant height and days to maturity (Kobo: 0.06; Meiso: −0.19; Sheraro: 0.01).

**Table 2.** Heritability and joint-linkage model power for each trait in the sorghum BC-NAM population

| Trait§ | Kobo |  |  | Meiso |  |  | Sheraro |  |  |
| --- | --- | --- | --- | --- | --- | --- | --- | --- | --- |
| | $H^2$¶ | No. QTL | Model $R^2$□ | $H^2$ | No. QTL | Model $R^2$ | $H^2$ | No. QTL | Model $R^2$ |
| DF | 0.33 | 10 | 0.26 (79%) | 0.30 | 10 | 0.25 (83%) | 0.71 | 7 | 0.20 (28%) |
| DM | 0.34 | 4 | 0.12 (35%) | 0.27 | 5 | 0.12 (44%) | 0.25 | 6 | 0.15 (60%) |
| PH | 0.55 | 1 | 0.13 (24%) | 0.39 | 8 | 0.20 (51%) | 0.75 | 6 | 0.16 (21%) |

§DF: days to first flowering; DM: days to maturity; PH: plant height.
¶Broad-sense heritability of BC-NAM population.
□Variance explained by joint-linkage model after fitting family term and detected QTL.
Numbers in parentheses represent proportion of genetic variance explained by the joint-linkage model, which was calculated by dividing model $R^2$ by $H^2$ .

### Genetic dissection of adaptive traits

Association analyses for the three measured traits revealed 27, 15, and 15 QTLs across the three environments using the joint-linkage (JL) model for days to flowering, days to maturity, and plant height, respectively (Tables 2, 3). Among the 27 JL QTLs for days to flowering, 25 (92.6%) showed overlapping confidence interval with previous QTLs detected for the same trait from multiple studies in the Sorghum QTL Atlas database [Mace *et al*. (2019)] (Dataset S4). Similarly, 12 of the 15 JL QTLs for days to maturity (80.0%) and 13 of the 15 JL QTLs (86.7%) for plant height overlapped with those found in previous studies (Dataset S4). Despite this validation of most QTLs, the total phenotypic variance explained by the final JL model was generally low, ranging from 0.20 for plant height to 0.26 for days to flowering in Kobo, from 0.12 for days to maturity to 0.15 for plant height in Meiso, and from 0.13 for days to flowering to 0.20 for days to maturity in Sheraro (Table 2). Heritability imposes an approximate upper limit to the *R*^2^ of a QTL model (Yu *et al*. 2008). Therefore, this is not unexpected given the low to moderate heritabilities of these adaptive traits under the natural drought conditions in Ethiopia. By taking the broad-sense heritability into account, proportions of genetic variance explained by the final joint-linkage models of each trait ranged from 28% to 83% in Kobo, 35%-60% in Meiso, and 21-51% in Sheraro (Table 2).

**Table 3.**
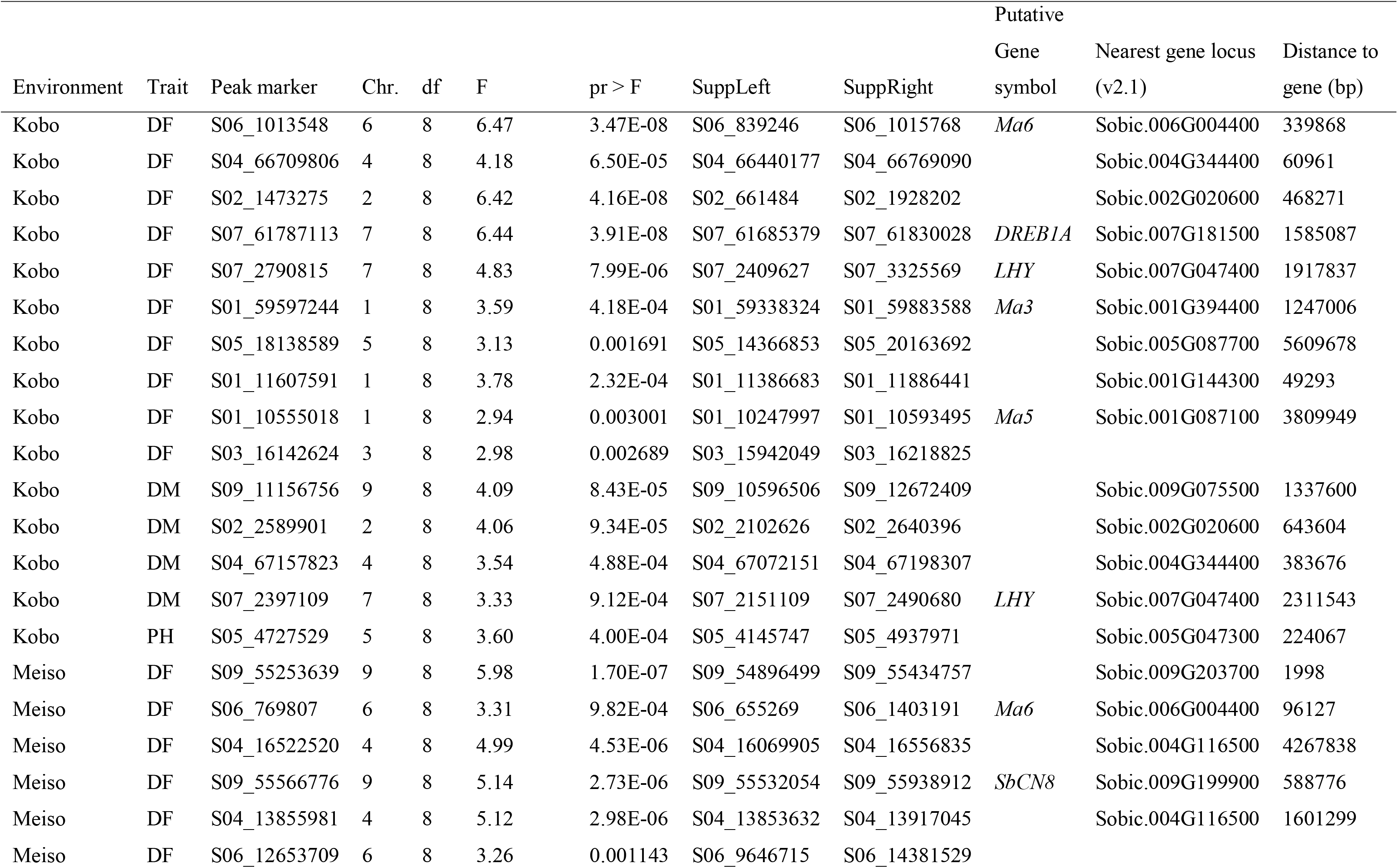
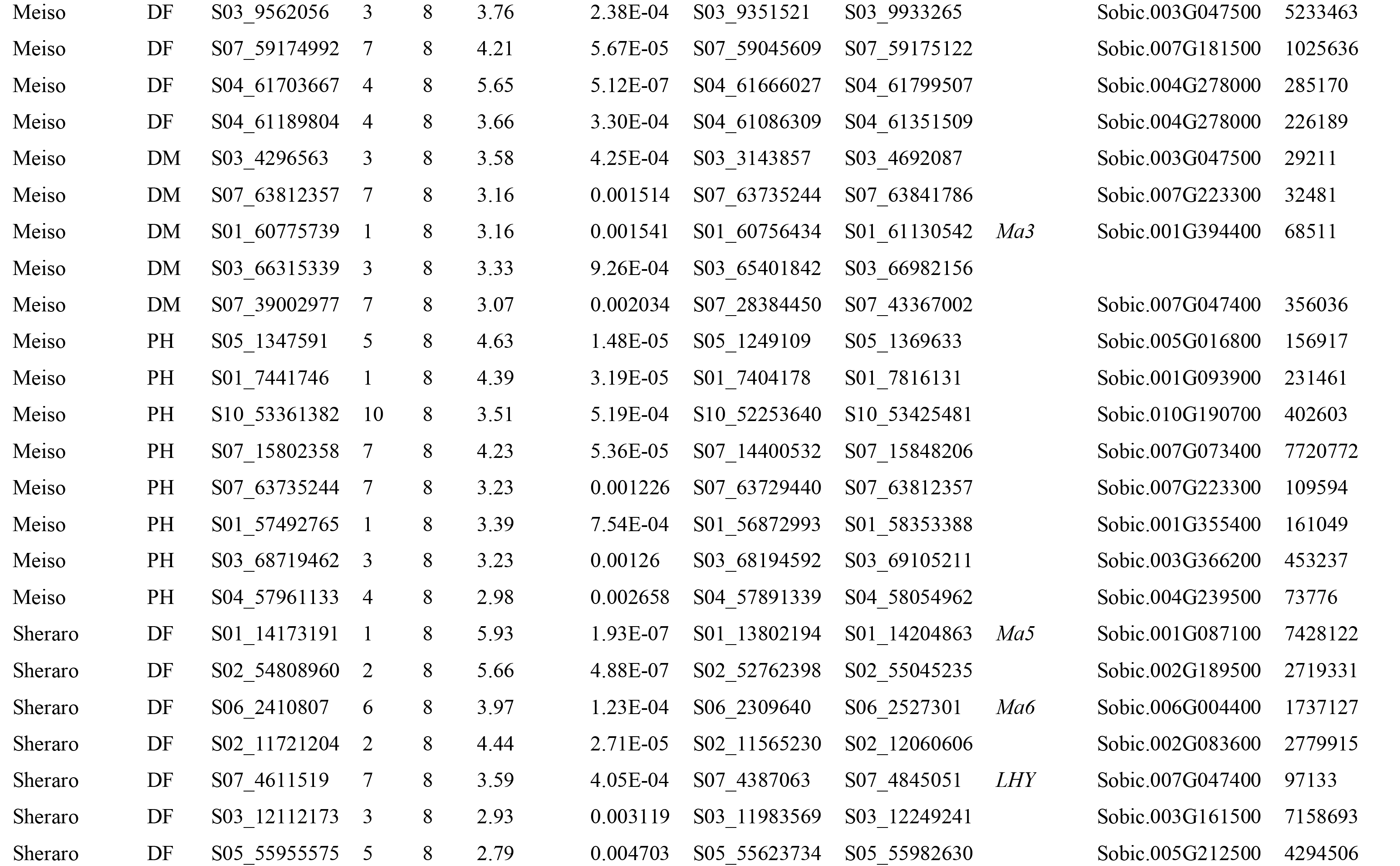
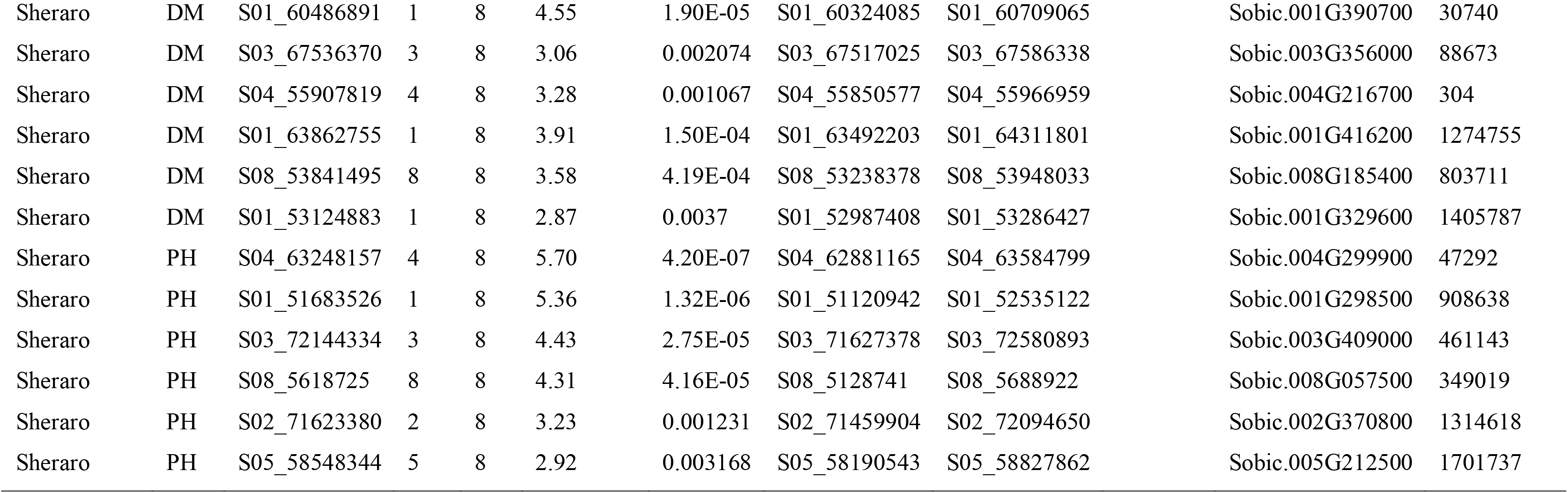
Summary of joint-linkage QTLs for days to flowering, days to maturity, and plant height in the sorghum BC-NAM population within three environments in Ethiopia.

In order to leverage the investment in their analysis, we incorporated the four small populations (IS2205, IS14556, IS16044, and IS32234) into GWAS analyses. The GWAS model included the same fixed effect of family term as the JL model, but marker effects were not nested within family. A total of 43, 6, and 35 SNPs exceeded the 10E-3 threshold for days to flowering, days to maturity, and plant height across three environments, respectively, which corresponded to 19, 4, and 15 likelihood peaks (Figures 5, 6, S2, Dataset S5). Despite the different statistical frameworks, there was generally high correspondence between JL QTLs and GWAS signals (Figures 5, 6, S2). Exact overlap of linkage mapping and GWAS is not expected, as linkage mapping tests markers within an individual population whereas GWAS tests marker effects across populations, with different strengths and weaknesses of each approach (Tian *et al*. 2011). Joint-linkage analysis produces many more small effects than GWAS analysis as an artifact of the model fitting process, which assigns a separate effect to all populations at each QTL. Moreover, the addition of four small populations in GWAS analyses may also confer discrepancies between these two methods.

**Figure 5.**
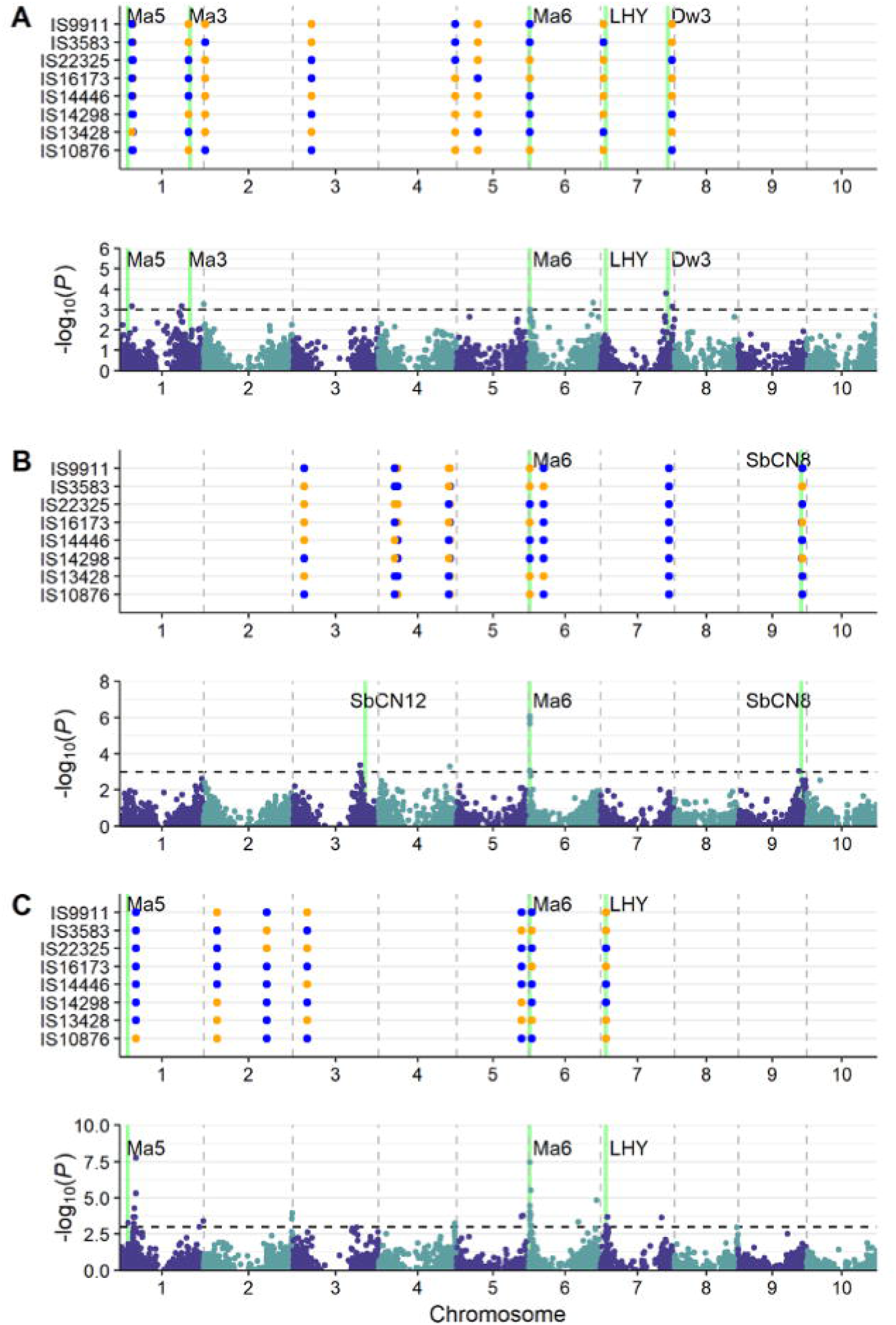
Marker-trait associations for days to flowering in (A) Kobo, (B) Meiso, and (C) Sheraro. Each panel shows associations detected in joint-linkage (top) and GWAS (bottom) models. Candidate genes were shown in green vertical lines and annotated with gene names.

**Figure 6.**
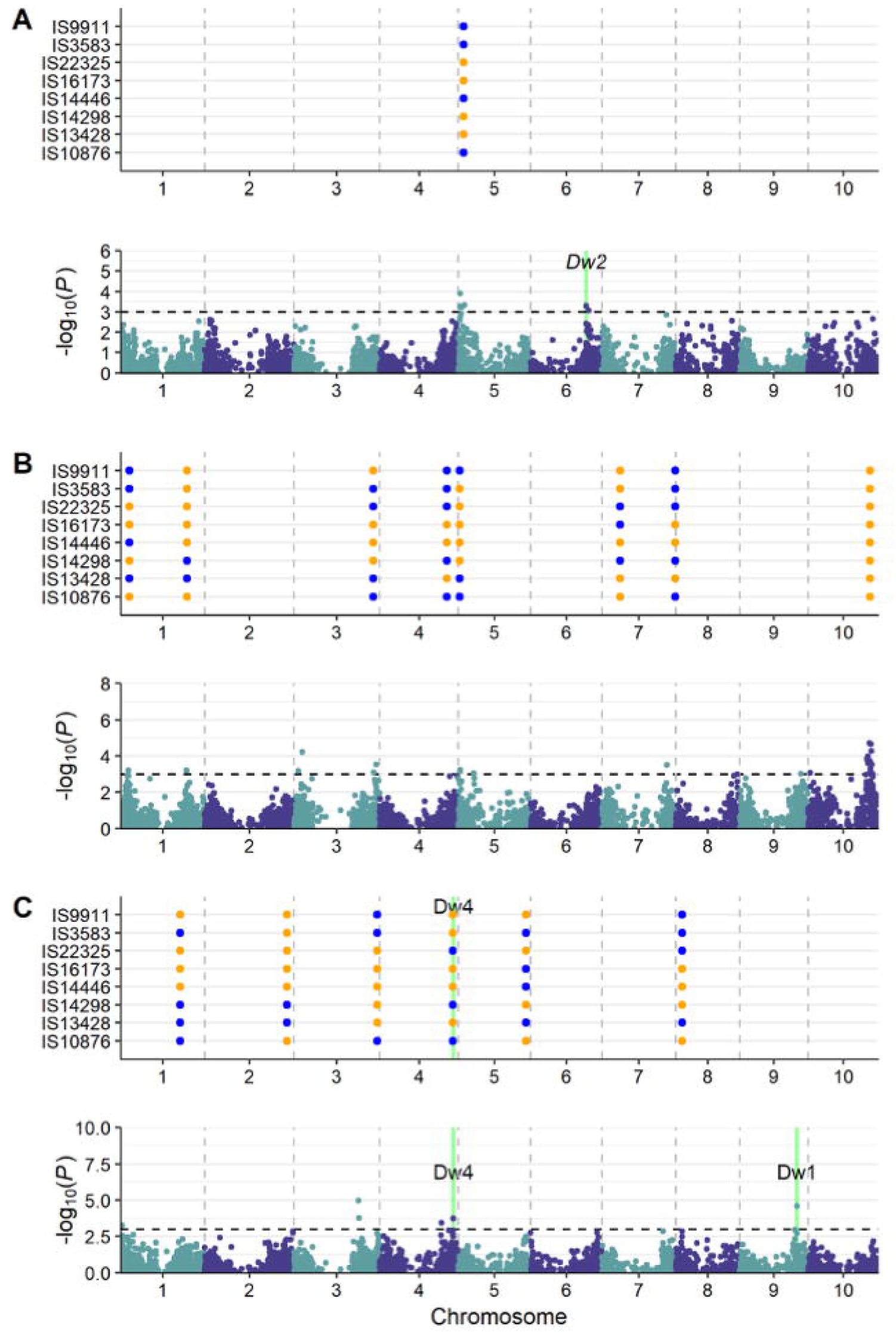
Marker-trait associations for plant height in (A) Kobo, (B) Meiso, and (C) Sheraro. Each panel shows associations detected in joint-linkage (top) and GWAS (bottom) models. Candidate genes were shown in green vertical lines and annotated with gene names.

Flowering time is one of the most important adaptive traits in grasses. The JL model detected 10, 10, and 7 JL QTLs for days to flowering at Kobo, Meiso, and Sheraro, respectively (Tables 2, 3, Figure 5), explaining 79%, 83%, and 28% of genetic variance (Table 2). Several QTLs were consistently detected across three environments near known sorghum maturity genes (Figure 5). The most significant QTL for days to flowering was detected at the putative *Ma6* gene (*CONSTANS*-like 4; Sobic.006G004400) on chromosome 6. In the JL analysis, the QTL peaks for days to flowering were 96 kb (S06_769807), 339 kb (S06_1013548), and 1.7 Mb (S06_2410807) from a candidate *Ma6* gene [*SbGHD7*, (Murphy *et al*. 2014)] in Meiso, Kobo, and Sheraro, respectively (Table 3, Figure 5). The GWAS model also consistently detected associations adjacent to *Ma6* in Kobo (S06_1015768), Meiso (S06_769807), and Sheraro (S06_769807) at 342 kb, 96 kb, and 96 kb away from *SbGHD7*, respectively (Figure 5, Dataset S5). One JL QTL peak (S09_55566776) in Meiso was detected about 588 kb downstream of the *SbCN8* gene (Sobic.009G199900), which encodes phosphatidylethanolamine-binding protein (PEBP) and is an ortholog of maize *ZCN8* and rice *OsFTL10*. A marginally significant GWAS association (S09_52569648) was also detected near *SbCN8* (Figure 5B, Dataset S4). In Meiso, the GWAS model also detected an association peak (S03_58246694) near *SbCN12* but the JL model did not detect signals near this region (Figure 5B). The *LHY* gene (LATE ELONGATED HYPOCOTYL; Sobic.007G047400) is 97 kb from a JL QTL (S07_4611519) in Sheraro on chromosome 7 (Figure 5C, Dataset S4) and 1.9 Mb from a JL QTL in Kobo (S07_2790815) that was (Figure 5A, Table 3). Strong associations with flowering were also detected on sorghum chromosome 1 at Kobo and Sheraro, which were relatively distant (more than 3 Mb) from the *Ma5* gene (Sobic.001G087100). Associations on chromosomes 2, 3, and 4 (Figure 5, Table 3, Files S4, S5) may represent novel genes.

For days to maturity, the JL model detected 4, 5, and 6 QTLs at Kobo, Meiso, and Sheraro, respectively, explaining 35%, 44%, and 60% of genetic variation (Table 2); while the GWAS model detected 2, 1, and 1 weak peaks (Figure S2). Days to flowering QTLs were mostly not near canonical maturity genes, except for one GWAS hit (S06_1015768) in Meiso near *Ma6* (Figure S2, Dataset S5). The majority of GWAS hits for days to maturity were only marginally significant (Figure S2, Dataset S5), perhaps due to the very limited phenotypic variation for this trait (Figure 4).

For plant height, the JL model detected 1, 8, and 6 QTLs at Kobo, Meiso, and Sheraro, respectively, explaining 24%, 51%, and 21% genetic variation (Figure 6, Tables 2, 3). The GWAS model detected 2, 8, and 5 peaks at these three environments (Figure 6, Dataset S5). Several JL QTLs and GWAS hits were adjacent to known sorghum dwarfing candidate genes. One GWAS hit (S06_48457872) from Kobo was approximately 6 Mb from the *Dw2* candidate gene (Sobic.006G067700) (Dataset S5), which is suggested to encode a protein kinase (Hilley *et al*. 2017). One JL QTL (S04_63248157) and GWAS hit (S04_63653942) at Sheraro were detected in the vicinity of the *Dw4* candidate gene, thought to be near 66.7 Mb on chromosome 4 (Li *et al*., 2015). An additional GWAS hit (S09_49635018) was among the most significant associations for plant height in Sheraro but was far from the *Dw1* candidate gene (Sobic.009G229800).

QTL allele effects that deviate from the prediction of parental phenotypic values indicate opportunities for selecting ‘transgressive’ progeny with values that exceed those of the more extreme parent. For days to flowering, 27 JL QTLs detected in the eight large populations (Tables 2, 3), permit estimation of a total of 216 (*i.e*. 27 × 8) QTL allelic effects – among these, 102 were negative (*i.e*. with the donor allele conferring earlier flowering than Teshale) and 114 were positive (*i.e*. with the donor allele conferring later flowering than Teshale) (Figure 7, Dataset S4). QTL allelic effects ranged from −7.4 to 5.1 days. At 26 of these 27 QTLs, both positive and negative alleles from donor parents were observed (Figure 5, Dataset S4), except for one QTL (S07_59174992) detected in Meiso at which all donor parents contributed positive effect alleles (Figure 5B, Dataset S4). On the other hand, across the 27 days to flowering QTLs, each donor parent contributed at least one positive and one negative allele(s). For days to maturity, the 120 QTL allelic effects of 15 JL QTLs (Tables 2, 3) included 61 negative (*i.e*. early maturity) and 59 positive effects (*i.e*. late maturity), ranging from −2.3 to 2.5 days, again with both positive and negative allelic effects at each QTL and from each donor (Figure S2, Dataset S4). For 120 QTL allelic effects estimated for the 15 JL plant height QTLs with a range from −23.8 cm to 18.1 cm (Figure 6, Dataset S4), fifty were positive (*i.e*. tall stature) and 70 negative (*i.e*. short stature). Similar to the other two traits, all plant height QTLs contained both positive and negative effects except for one QTL (S10_53361382) in Meiso, where all donor parents contributed negative effect alleles.

**Figure 7.**
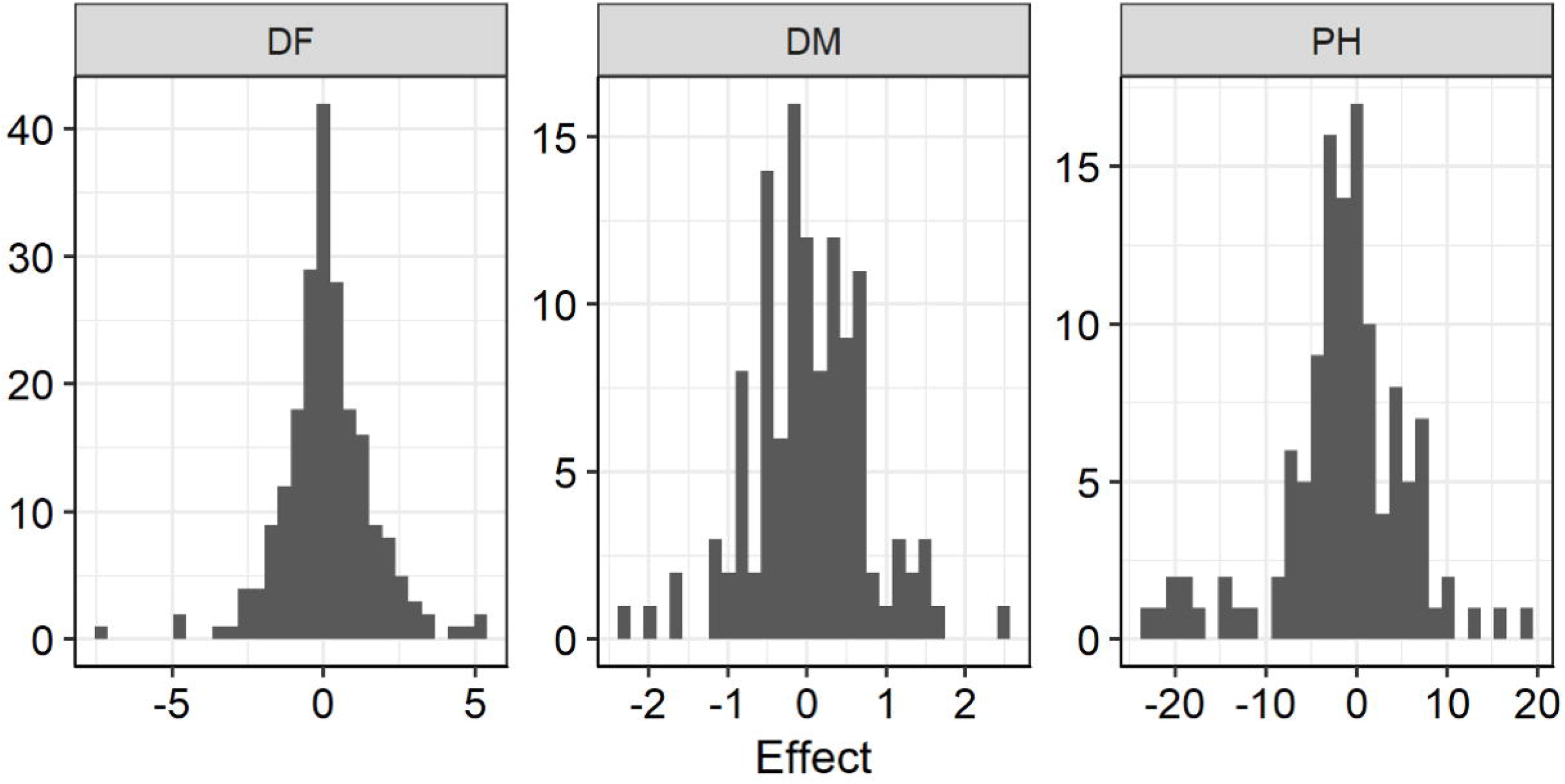
Distributions of allelic effects of joint-linkage QTLs detected in a sorghum BC-NAM population. Inside each plot, x-axis represents allelic effects and y-axis represents frequency. DF: days to 50% flowering, DM: days to maturity, PH: plant height.

## Discussion

A backcross nested association mapping population made by crossing thirteen diverse sorghum lines to an elite cultivar (Teshale) bred for the center of diversity (Ethiopia) provides insight into the biogeography of trait variation. The backcross nested breeding design combines practical breeding efforts for introgression of new alleles into adapted germplasm with statistical power to dissect quantitative traits (Buckler *et al*. 2009; Jordan *et al*. 2011; Yu *et al*. 2008). Employing donor lines chosen for divergent drought defense responses, this BC-NAM population also increases the genetic diversity available in Ethiopian elite adapted sorghum germplasm, providing new scope to improve food security of societies dependent upon this crop in a region known for periodic devastating droughts.

Multiple genomic properties attest to the usefulness of this sorghum BC-NAM population (Figures 2, 3, Table 1). Principal component analysis displayed clear population structure, indicating minimal cross contamination among families (Figure 2A). Indeed, most individual populations clustered together except for IS22325 and IS14298. This was likely due to the backcross breeding scheme used in this study. Molecular marker analysis indicated retention of an average of 76.3% of the recurrent parent genome in this BC-NAM population, close to the expected 75% (Figure 2B).

This BC-NAM population was effective in dissecting quantitative traits. Our study identified 27, 15, and 15 QTLs for days to flowering, days to maturity, and plant height, respectively (Tables 2, 3). Both the present study and another of different germplasm [Mace *et al*. (2013)] found that genetic control of flowering time in sorghum is substantially more complex than classical genetics was able to resolve (Quinby 1974), involving a relatively large number of loci with small effects, as we have suggested (Zhang *et al*. 2015). A degree of validation is provided by the observation that many of the detected QTLs in these two BC-NAM populations overlapped with those detected by multiple bi-parental mapping studies (Dataset S4).

The ability to resolve many QTLs together with the ability to sample more allelic diversity than bi-parental populations reveals the spectrum of allele effects in the study population, in comparison to those of the common parent. In this case, the common parent, Teshale, is strategically chosen in that it was bred in and selected for a target environment near the species center of diversity – while the other parents were selected from a broad sampling of germplasm based on their diverse spectrum of drought responsiveness traits (Vadez *et al*. 2011). The finding that for all three measured traits, nearly equal numbers of alleles conferred increases and decreases in phenotype relative to the Teshale allele, is consistent with the notion that Teshale is well adapted to the center of diversity for this particular gene pool, presumably with a history of balancing selection, while the 13 exotic sorghum lines from locales widely-distributed across the natural and introduced range sample smaller marginal populations in which novel alleles may be more likely to persist due to the effects of selection and genetic drift.

This work exemplifies the nature of efforts that may be necessary to adapt many crops to new climate extremes, with the introduction of novel or extreme traits from exotic germplasm necessitating a new epoch of selection to re-establish an adaptive peak or reach a new one. With a worldwide water crisis looming, developing drought tolerant sorghum is vital in rain-fed environments, particularly in sub-Saharan Africa. In Ethiopia, agriculture is the largest economic sector and contributes 48% of the nation’s GDP, and sorghum provides more than one third of the cereal diet and is widely grown for food, feed, brewing, and construction purposes (Ethiopian Institute of Agricultural Research 2014). However, sorghum yield in Ethiopian could decrease by 40% as intense climate change events become more common and droughts are likely to become more prevalent early in the growing season when crops are vulnerable (Eggen *et al*. 2019). Genetic diversity within breeding programs decreases due to selection, small population size, genetic drift and other factors (Fu 2015; Reif *et al*. 2005) and reaching outside of local breeding programs will be necessary to adapt many crops to new climate extremes.

In a companion paper (Dong et al. unpublished), phenotypic performance of this BC-NAM population under drought environments showed that the drought resilience of Teshale can be improved by incorporation of different adaptations from the diverse donor lines, however none of the resulting populations produced higher population mean yield than Teshale – meaning that to arrive at a new ‘adaptive peak’ (Dong et al. unpublished), selection to traverse a ‘valley’ of reduced yield is necessary. Selection response for quantitative traits is determined by genetic variance, heritability, and selection intensity (Falconer & Mackay 1996). Rich variation reflected by mixtures of ‘positive’ and ‘negative’ QTL alleles for all traits provides a foundation for selection, with new diversity from the diverse donor lines complementing the adaptive phenotype of Teshale. The finding in our companion paper (Dong et al. unpublished) of correlations of plot-based grain yield with days to flowering (−0.20 to −0.42); and plant height (0.14-0.39) exemplify scope for the sorts of adjustments that may be needed to re-establish locally adaptive phenotypes. Indeed, with the enormous altitudinal variation of a country such as Ethiopia, somewhat different lines may be needed for different locales.

## Acknowledgments

This study was supported by United States Agency for International Development (USAID) awarded to AHP (grant no.: AID-OAA-A-13-00044). The authors thank Jimma University, Ethiopian Institute of Agricultural Research, Amhara Region Agricultural Research Institute and Tigray Region Agricultural Research Institute for providing land and technical support for the field trials. The authors also thank the members of the Paterson Lab for valuable help and discussions. We also thank Georgia Genomics and Bioinformatics Core for sequencing service.

## Author Contribution

AHP and KB jointly developed and led this project. TB and KB developed the populations, TB, A, MW, and KB conducted field trials, and phenotypic data collection. HD and CL made GBS libraries. HD performed data analysis and wrote the draft manuscript. All authors commented and reviewed the manuscript.

