## Supplementary material for "New adaptive peaks for crops – an example from improvement of drought-resilience of sorghum in Ethiopia": Figure S1, Figure S2, Table S1, Table S2, Table S3

### *Plants, People, Planet* Supporting Information

The following Supporting Information is available for this article:

**Fig. S1** Principal component analysis (PCA) within each population. Suspected contaminants highlighted in red were discarded.

**Fig. S2** Marker-trait associations for days to maturity in (A) Kobo, (B) Meiso, and (C) Sheraro. Each panel shows associations detected in joint-linkage (top) and GWAS (bottom) models. Candidate genes were shown in green vertical lines and annotated with gene names

**Table S1** Number of BCNAM lines survived in each population within each environment.

| Environment | N_1 rep_^¶^ | N_2 rep_^§^ |
| --- | --- | --- |
| Kobo | 362 (58) | 715 |
| Meiso | 195 (58) | 1017 |
| Sheraro | 2 | 1072 |

¶No. of BCNAM lines with only one replicate survived. Number in parenthesis represent number of lines that lost one replicate in both Kobo and Meiso.

§No. of BCNAM lines with both replicates survived.

**Table S2**. Analysis of variance for three traits studied in this sorghum BCNAM population. This population was tested at three locations in Ethiopia: Kobo, Meiso, and Sheraro.

| Trait§ | Model term¶ | DF | SS | MS | F | Pr (>F) |
| --- | --- | --- | --- | --- | --- | --- |
| DF | Genotype | 1086 | 66748 | 61 | 2.15 | <2.2e-16 |
|  | Location | 2 | 409240 | 204620 | 7155.25 | <2.2e-16 |
|  | Genotype * Location | 2004 | 87085 | 43 | 1.52 | <2.2e-16 |
|  | Residual | 2602 | 74410 | 29 |  |  |
| DM | Genotype | 1086 | 85402 | 79 | 3.52 | <2.2e-16 |
|  | Location | 2 | 1547147 | 773573 | 34658.69 | <2.2e-16 |
|  | Genotype * Location | 2004 | 57708 | 29 | 1.29 | <5.7e-10 |
|  | Residual | 2602 | 58076 | 22 |  |  |
| PH | Genotype | 1086 | 1601184 | 1474 | 3.97 | <2.2e-16 |
|  | Location | 2 | 13648406 | 6824203 | 18373 | <2.2e-16 |
|  | Genotype * Location | 2004 | 1404901 | 701 | 1.89 | <2.2e-16 |
|  | Residual | 2602 |  |  |  |  |

§DF: days to 50% flowering, DM: days to maturity, PH: plant height

¶Note that for the multi-environment ANOVA, only the nine populations (IS10876, IS15428, IS14298, IS14446, IS16173, IS22325, IS23988, IS3583, IS9911) that included at all three locations were used. In terms of model terms, initial model also included population, and genotype nested in population, however, population term and genotype nested in population term ran into ‘perfect multicollinearity’ problem, and thus population term was removed from ANOVA model.

**Table S3**. Sorghum BCNAM trait means within each environment.

| Trait§ | Unit | Kobo | Meiso | Sheraro |
| --- | --- | --- | --- | --- |
| DF | days | 78.15 | 85.06 | 64.99 |
| DM | days | 127.04 | 122.67 | 90.22 |
| PH | cm | 175.66 | 177.72 | 279.60 |

§DF: days to 50% flowering, DM: days to maturity, PH: plant height

**Dataset S1** Phenotypic data summary

**Dataset S2** SNP information

**Dataset S3** BC-NAM lines information

**Dataset S4** Summary of joint-linkage QTL

**Dataset S5** Summary of GWAS hits
